# scGate: marker-based purification of cell types from heterogeneous single-cell RNA-seq datasets

**DOI:** 10.1101/2021.11.08.467740

**Authors:** Massimo Andreatta, Ariel J. Berenstein, Santiago J. Carmona

**Affiliations:** Ludwig Institute for Cancer Research, Lausanne Branch, and Department of Oncology, CHUV and University of Lausanne, Epalinges 1066, Switzerland; Swiss Institute of Bioinformatics, Lausanne, Switzerland; Laboratorio de Biología Molecular, División Patología, Instituto Multidisciplinario de Investigaciones en Patologías Pediátricas (IMIPP), CONICET-GCBA, Buenos Aires C1425EFD, Argentina

## Abstract

A common bioinformatics task in single-cell data analysis is to purify a cell type or cell population of interest from heterogeneous datasets. Here we present scGate, an algorithm that automatizes marker-based purification of specific cell populations, without requiring training data or reference gene expression profiles. scGate purifies a cell population of interest using a set of markers organized in a hierarchical structure, akin to gating strategies employed in flow cytometry. In our benchmark for blood-derived and tumor-infiltrating immune cells, scGate outperforms SingleR, a state-of-the-art classifier for single-cell data. scGate is implemented as an R package and integrated with the Seurat framework, providing an intuitive tool to isolate cell populations of interest from complex scRNA-seq datasets.

**Availability:** R package source code and reproducible tutorials are available at https://github.com/carmonalab/scGate

## Introduction

Single-cell RNA sequencing (scRNA-seq) is becoming increasingly popular, enabling high-throughput exploration of cell types and states from complex tissues. Cell types are generally defined based on biological function and the markers used to physically isolate them, but these can change depending on the source tissue. In scRNA-seq data analysis, knowledge of cell type-defining marker genes is typically used to manually identify relevant cell populations within custom bioinformatics workflows, requiring several steps and parameters.

Alternatively, when high-quality transcriptomic profiles are available for the cell type of interest, training multinomial machine learning classifiers to predict cell type identity has been shown to be a powerful approach [1,2]. For example, popular tools such as SingleR perform well when trained on high-quality bulk RNA-seq gene expression profiles of sorted cell populations [2]. However, reliable reference transcriptional profiles are not always available. Moreover, batch effects and other biases are difficult to assess in training datasets, which can lead to overfitting and biased predictions.

In this work, we developed an intuitive tool to purify a cell population of interest from complex scRNA-seq datasets based on literature-derived marker genes, without requiring reference gene expression profiles or training data. With scGate, an expert can purify a cell population of interest from a complex scRNA-seq dataset by only defining a few marker genes, or by using sets of markers provided with the scGate package. This provides a straightforward and complementary approach to machine learning-based classifiers, aimed at automatizing current practices in marker-based purification of cell types from single-cell transcriptomics datasets.

## Results

We developed scGate, an R package that automatizes the typically manual task of marker-based cell type annotation, to enable accurate and intuitive purification of a cell population of interest from a complex scRNA-seq dataset (for instance, a dataset of blood-derived immune cells, Figure 1A). scGate builds upon our recent method UCell [3] for robust single-cell signature scoring, and Seurat, a comprehensive and powerful framework for single-cell omics analysis [4].

**Figure 1.**
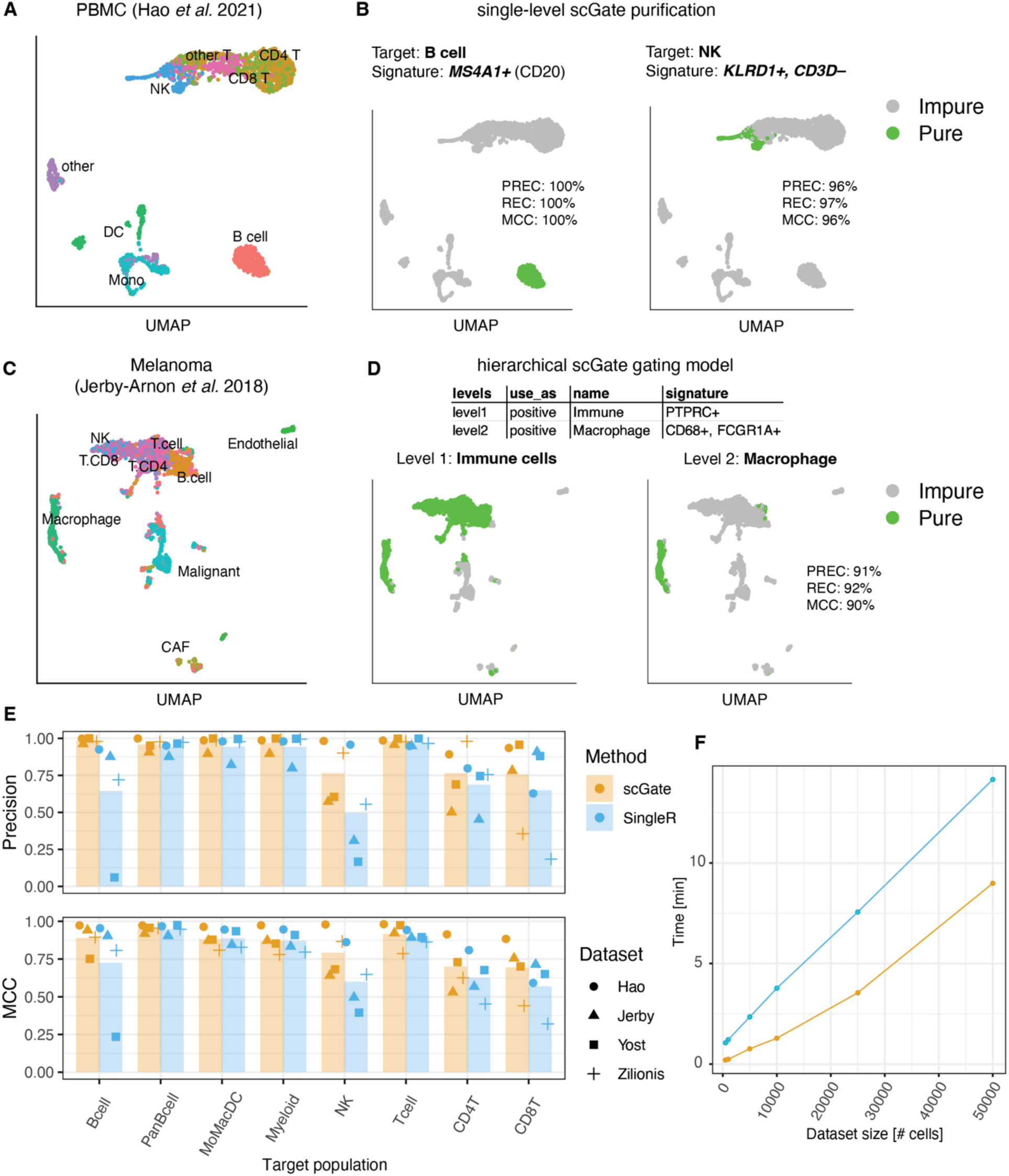
Purifying cell populations from single-cell datasets using scGate. **A)** Uniform Manifold Approximation and Projection (UMAP) representation of scRNA-seq data of peripheral blood mononuclear cell (PBMC) populations annotated by Hao et al. **B)** Purification of target cell types using scGate, for B cells on the left (using marker CD20 encoded by *MS4A1*) and natural killers (NK) on the right (using *KLRD1* positive and *CD3D* negative markers); precision (PREC), recall (REC) and classification accuracy (MCC) are shown. **C)** UMAP representation of scRNA-seq data of melanoma tumors annotated by Jerby-Arnon et al. **D)** Purification of macrophages using a hierarchical gating model: immune cells at the first level (left panel) and macrophages at the second level (right panel). **E)** Precision (Positive Predictive Value) and MCC (Matthews Correlation Coefficient) values for 4 publicly available scRNA-seq datasets (derived from blood or tumors) for scGate and SingleR, trained using recommended reference profiles. **F**) Running time as a function of dataset size for scGate (using a complex 4-level GM) and SingleR (using HPCA database) on a 16GB RAM machine, using 4 cores.

Briefly, scGate takes as input: *i)* a gene expression matrix or Seurat object and *ii)* a “gating model” (GM), consisting of a set of marker genes that define the cell population of interest. The GM can be as simple as a single marker gene, or a combination of positive and negative markers. For example, the marker *MS4A1* (encoding CD20) alone allows purifying B cells with 100% precision and 100% recall (Figure 1 B left panel). A model that requires *KLRD1* but absence of *CD3D* purifies natural killer (NK) cells with 96% precision and 97% recall (Figure 1 B right panel). More complex GMs can be constructed in a hierarchical fashion. For instance, macrophages can be purified from a complex tissue such as melanoma tumors (Figure 1 C) by defining a two-level hierarchical gating model. The first level gates on immune cells using pan-immune cell marker *PTPRC* encoding CD45, and subsequently the second level purifies macrophages from immune cells using the markers *CD68* and *FCGR1A* (Figure 1 D).

Our algorithm evaluates the strength of the marker gene expression in each cell using the rank-based method UCell, and then performs k-nearest neighbor (kNN) smoothing by calculating the mean UCell score across neighboring cells. By kNN-smoothing, scGate aims at mitigating the large degree of sparsity in scRNA-seq data. Finally, a fixed threshold over kNN-smoothed signature scores is applied in binary decision trees generated from the user-provided gating model (e.g. Figure 1 D), to annotate cells as either “pure” or “impure” with respect to the cell population of interest.

The intuitive and flexible design of scGate allows for positive and negative markers and sequential/hierarchical gating strategies, providing users a quick and simple, yet powerful tool to purify cell populations of interest from arbitrarily complex datasets. Each of the purifications shown in Figure 1 B required just one line of R code within a Seurat workflow, for instance, to purify NK cells:

~~~
scGate(data = seurat_object, model = gating_model(name = “NK”, signature = c(“KLRD1”,“CD3D-”))
~~~

scGate comes with pre-defined GMs based on commonly used markers of immune cell types in human and mouse, such as T cells, B cells, NK cells, myeloid cell populations, among others. With these marker sets and four author-annotated published datasets from blood or tumors [4–7], we compared the predictive performance of scGate against one of the most popular and top-performing single-cell classifiers, SingleR [8]. Of note, SingleR requires reference gene expression profiles for training. In this case, we used the recommended “Human Primary Cell Atlas” (HPCA) dataset for training SingleR and other parameters by default. Across the board, scGate showed a superior mean precision (0.89 vs 0.78, paired wilcoxon test p-value = 1.8 10^−5^) and predictive power (Matthews Correlation Coefficient/MCC 0.84 vs 0.76, paired wilcoxon test p-value = 7.7 10^−4^) compared to SingleR (Figure 1 E), as well as a faster computing time (Figure 1 F).

Multiple predefined scGate models are provided in a public repository as version-controlled tab-separated text, allowing scGate to automatically synchronize its internal database of gating models. Users can manually edit these models and easily write their own. scGate also provides functions to evaluate the performance of custom gating models, either user-provided or those that accompany the package, on a set of pre-annotated “testing” datasets. Overall, scGate provides an accurate, scalable and intuitive tool to isolate cell populations of interest that can be seamlessly incorporated into Seurat pipelines for single-cell data analysis.

## Availability and Implementation

scGate is available as an R package at https://github.com/carmonalab/scGate. A reproducible workflow describing the main functions and usage of the package, as well as the code to reproduce the benchmark, can be found at https://github.com/carmonalab/scGate.demo and https://github.com/carmonalab/scGate.benchmark.

